# Genetic Association of Somatic Incompatibility and NLR-like Protein Domains in *Coprinopsis cinerea*

**DOI:** 10.64898/2026.06.24.733965

**Authors:** Ben Auxier, Julia Marschall, S. Lorena Ament-Velásquez, Johan Baars, Karin Scholtmeijer, Arend van Peer, Alfons J.M. Debets, Duur K. Aanen

## Abstract

In fungi, hyphal fusion is beneficial within an individual, but fusion between individuals comes with the risks of infection or exploitation. To manage this risk, fungi have developed mechanisms to restrict sustained fusion to be within a genetic individual, called allorecognition. In Ascomycete fungi, this recognition is based on allelic identity at several polymorphic allorecognition genes, often triggering cell death. However, the genetic basis of allorecognition is unknown in basidiomycetes, the clade that includes mushroom-forming fungi. Here, we map the first locus for this trait, which we call *somA*, in the mushroom-forming fungus *Coprinopsis cinerea*. We combined F1 offspring phenotypes with independent backcross lines to identify a region on chromosome 5 linked with the production of a barrage zone, a classic allorecognition phenotype. Fine-mapping of this region resulted in a region with a set of kinases and NACHT domain proteins, flanked by a leucine-rich repeat (LRR) protein. While the NACHT and kinase proteins are diverse between the parents, the LRR-encoding protein shows signs of purifying selection. Additional *C. cinerea* genomes show that this region contains several highly divergent alleles, consistent with long-term balancing selection. These polymorphic alleles all contain a single monomorphic LRR, which may indicate a novel mechanism for fungal nonself recognition. Based on a phylogenetic survey of related Basidiomycetes, this specific locus architecture appears to be restricted to closely related species. This finding of a multiallelic locus may explain the general trend of few nonself recognition loci in basidiomycetes. These results provide a first understanding of how individuality is maintained in basidiomycetes.

**Significance Statement:** How mushroom-forming fungi recognize each other as individuals is an open question. Here, we identify the first genes that trigger this recognition in a mushroom-forming fungus. As these fungi have very distinctive lifecycles compared to mold-forming fungi, it was hypothesized that the process would operate from different mechanisms. Our results show the molecular mechanisms are in fact quite similar. These results provide a first step towards understanding how these fungi can both fuse with themselves and still discriminate against other individuals.

## Introduction

The concept of individuality in basidiomycete fungi, the group that produces mushrooms as well as rusts and smuts, is difficult to define. These fungi exist both as monokaryons, composed of haploid nuclei with a common genotype, as well as dikaryons, where two distinct haploid genotypes stably coexist in a single mycelial network. Initially, mycologists thought that interindividual fusion formed cooperative chimeric networks, termed the “Unit Mycelium”, distributing resources for the group’s benefit (Buller, 1930). However, later work clarified that these fungi readily distinguish self from nonself, competing instead of cooperating. In both field and laboratory experiments, interactions between distinct isolates produced heavily melanized barrage zones, while interactions between clones of an isolate did not, leading to Rayner’s influential synthesis of the “Individualistic Mycelium” (Rayner, 1991; Rayner et al., 1984), later confirmed in other species (Stenlid, 1985).

These interactions have been termed somatic incompatibility (SI), as the two mycelial networks, the bodies of the fungi, are not compatible with each other (Worrall, 1997). The SI phenotype between individuals seems to be somewhat reduced when individuals are more closely related (Hansen et al., 1993; May, 1988), indicating a system of multiple genetic loci. Segregation ratios of the SI phenotype between progeny from a single individual shows segregation consistent with 3-4 loci in the forest pathogens *Heterobasidion annosum* (Hansen et al., 1993), and *Collybia fusipes* (Marcais et al., 2000), as well as the button mushroom, Agaricus bisporus (Scholtmeijer et al., 2025). The number of loci seems variable, as two loci were found in *Amylostereum areolatum*, a symbiont with wood wasps (van der Nest et al., 2008), two loci in *Serpula lacrymans*, an invasive fungus in human building materials (Kauserud et al., 2006), and potentially a single locus in the tree pathogen *Phellinus gilvus* (Rizzo et al., 1995). Only for a few species has the SI phenotype been mapped, and this only with coarse QTL consisting of large chromosomal segments (Lind et al., 2007; Scholtmeijer et al., 2025; van der Nest et al., 2009).

In mushroom-forming basidiomycetes, there are two historic model genetic systems. Originally used for identification of the loci controlling mating compatibility (Raper & Raper, 1966; Wendland et al., 1995), *Schizophyllum commune* has become a widely used organism to understand both the biochemistry and lifecycle of basidiomycetes (Ohm et al., 2010; Wessels et al., 1991). However, *S. commune* lacks an obvious nonself recognition phenotype (Nieuwenhuis et al., 2013) so despite its considerable genetic toolbox, it is not suitable for genetic dissection of this trait. The other main genetic model in mushroom-forming fungi is *Coprinopsis cinerea*, which also has a well-defined sexual cycle, a high-quality annotated reference genome available (Moore & Pukkila, 1985; Stajich et al., 2010), and wild collections (M. Wu et al., 1983). Importantly, SI is well described in C. cinerea, where isolates from across the globe show SI response in nonself interactions consisting of a pigmented line between individuals (May, 1988).

While the molecular basis for SI in basidiomycetes is unknown, in ascomycete fungi, the sister phylum to basidiomycetes, the genes involved are often related to NOD-like receptors (NLRs), which are described in immune systems of both plants and animals and bacteria (Gao et al., 2022; Uehling et al., 2017). Proteins in the NLR family are characterized by an N-terminal effector domain, a central nucleotide binding domain, and C-terminal repetitive domain (Hu & Chai, 2023). In fungal nonself recognition genes, this central domain is often of the NACHT family (Koonin & Aravind, 2000). In the best-studied species *Neurospora crassa* and *Podospora anserina*, the HET protein domain is sometimes also found (Glass & Dementhon, 2006). This domain is found across fungal genomes, but in other groups of fungi does not appear to be associated with nonself recognition (Arshed et al., 2023; Choi et al., 2012). In an attempt to transfer knowledge from these model systems, a bioinformatic survey of basidiomycete genomes for candidate genes, based on *N. crassa* and *P. anserina*, pointed to some shared genes, but an overall absence of HET domains in basidiomycetes (van der Nest et al., 2014). Further investigations of NLR proteins in fungi found no clear candidates for what could be controlling basidiomycete SI (Dyrka et al., 2014).

Although the molecular components are variable across organisms, a common factor of genes involved in nonself recognition is balancing selection. As the nonself recognition alleles are most useful when rare, selection on these alleles becomes negative frequency-dependent (Spurgin & Richardson, 2010). This leads to alleles that are older than would be expected under neutral processes, often older than the species itself, resulting in alleles shared across species boundaries in trans-species polymorphisms (Charlesworth, 2006). Across plants, animals and fungi, the effects of balancing selection can be seen in population level DNA sequence data (Auxier et al., 2024; Hedrick & Thomson, 1983; Tian et al., 2002; J. Wu et al., 1998). The even allele frequencies and trans-species polymorphisms provide clues that balancing selection is acting. However, there can be multiple overlapping causes for this pattern of genetic diversity (Lobkovsky et al., 2019; Spurgin & Richardson, 2010).

The extensive research of ascomycete nonself recognition provides a background for understanding basidiomycete SI. However, bioinformatics surveys have failed to identify clear causal genes for this biologically important phenotype. Here we leverage the increased availability of sequencing information, using both genetic and bioinformatic techniques, to map genetic variants of SI in the model species *C. cinerea*. We identify a first locus, *somA*, associated with this trait. This complex locus contains hallmarks of nonself recognition genes in other fungi, but with a distinctive organization.

## Results

Interactions between dikaryons of Oko-7+FGSC#25194 and Java-6+FGSC#25194 (Fig. 1A) produced clear barrage reactions after four weeks which we did not observe in control self-pairing reactions (Fig. 1B). To genetically dissect this trait, we phenotyped 576 offspring of Oko-7 x Java-6, recovering a range of phenotypes, which we classified as 71 (12.3%) compatible to Okayama-7, and the rest as incompatible. The incompatibility phenotype ranged from dark pigmented lines to less visible gaps in the hyphal growth front between two interacting colonies (Fig. 1B).

**Figure 1:**
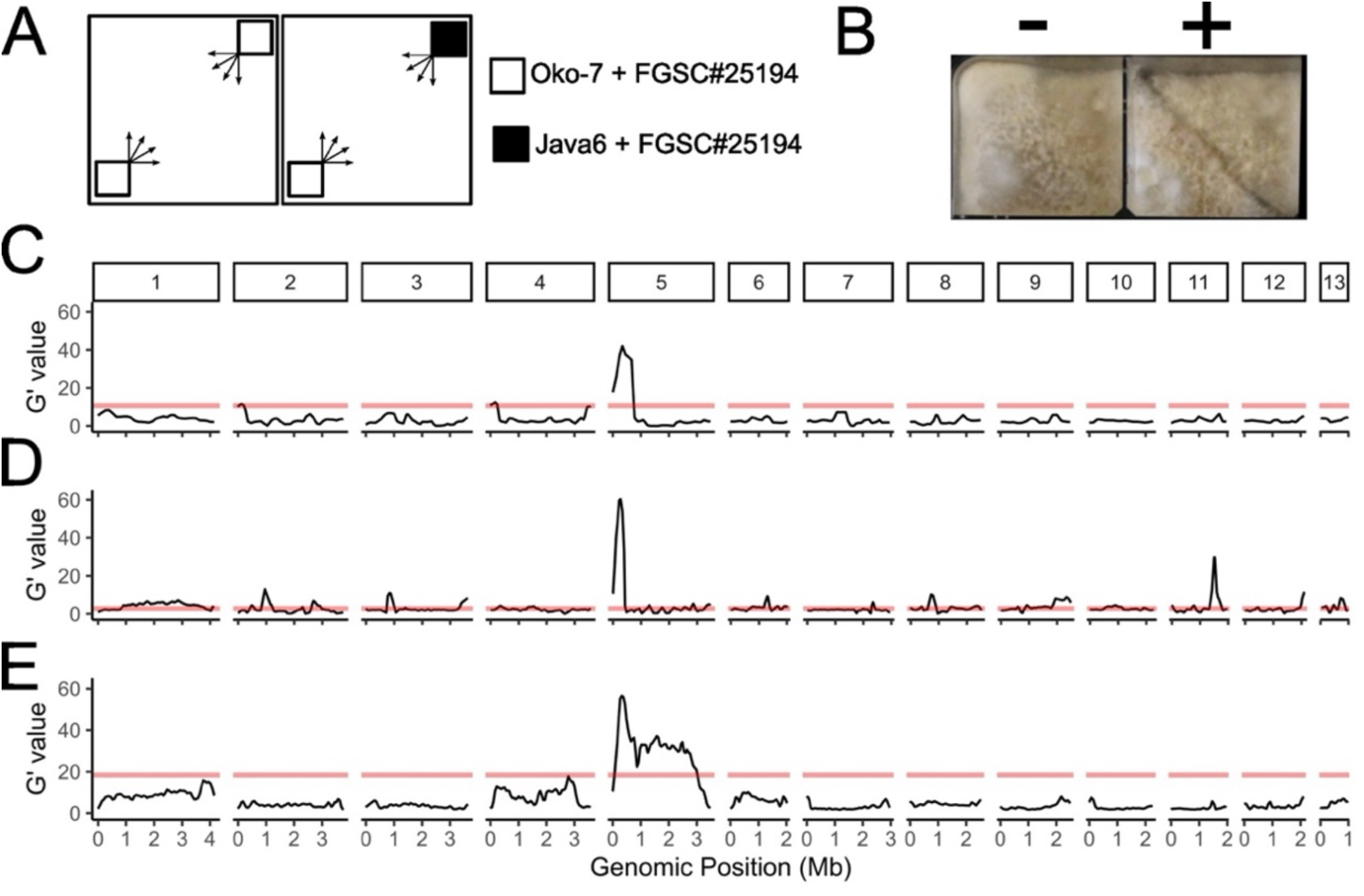
Genome-wide mapping of the somatic incompatibility response in *C. cinereus*. A) Schematic of interactions between mated dikaryons. B) Output of control reactions, self-pairing and positive control, that either produce no visible line, or a dark pigmented line in the interaction zone. C-E) Genetic mapping of somatic incompatibility in *C. cinerea*. G’ value represents difference in allele frequency between Compatible and Incompatible bulked groups smoothed over 100 kb windows across the 13 chromosomes. C) represents compatible and incompatible bulks from backcrossed line 11-13-8. D) bulks from backcrossed line 8-24-15-7. E) values from bulks of F1 progeny. Horizontal red lines indicate significant at α=0.01 based on a permutation test.

To genetically map this trait, we used bulk segregant analysis (BSA) based on 307,611 segregating variants we identified between Java-6 and the reference Okayama-7. Analysis of the compatible and incompatible F1 offspring bulks (65 compatible and 35 incompatible) showed a single peak located on the left-hand side of chromosome 5, with a G’ value of approximately 55, although the majority of the chromosome rose above the significant threshold (Fig. 1E). To reduce the genetic complexity, we backcrossed two lines of F1 offspring repeatedly to Okayama-7, selecting for somatically incompatibility, but sexual compatibility, at each round. BSA of pools of 20 compatible and 20 incompatible offspring from the 11-13-8 line (3x backcrossed) showed a single peak on chromosome 5, with variation restricted to a region spanning ∼0.5 Mb (Fig. 1C). A peak at a similar location was found in the BSA of 19 compatible and 51 incompatible offspring from the backcrossed line 8-24-15-7 (4x backcrossed) (Fig. 1D). The 8-24-15-7 backcrossed line also showed a second, smaller, peak on the right side of chromosome 11, albeit with a reduced genetic association (G’ ∼25). Being found in all three mappings, the region on the left side of chromosome 5 was selected for further study and termed *somA*, referring to its association with somatic incompatibility.

To fine map the *somA* region, we used progeny from the 8-24-15-7 backcrossed line crossed with Okayama-7. From 200 progeny, we recovered 47 recombinants across the region spanning 80 kb to 400kb of chromosome 5. Phenotyping of these offspring as well as genotyping at six markers refined the QTL peak to be between 200 kb and 300 kb, peaking at 275 kb (Fig. 2A). This peak association aligned with a drop in sequencing depth of Java-6 mapped against Okayama-7 (Fig. 2B). The window spanning 200-300 kb along Chromosome 5 contains 41 genes, with 20 genes in the narrower region of 250-300 kb.

**Figure 2:**
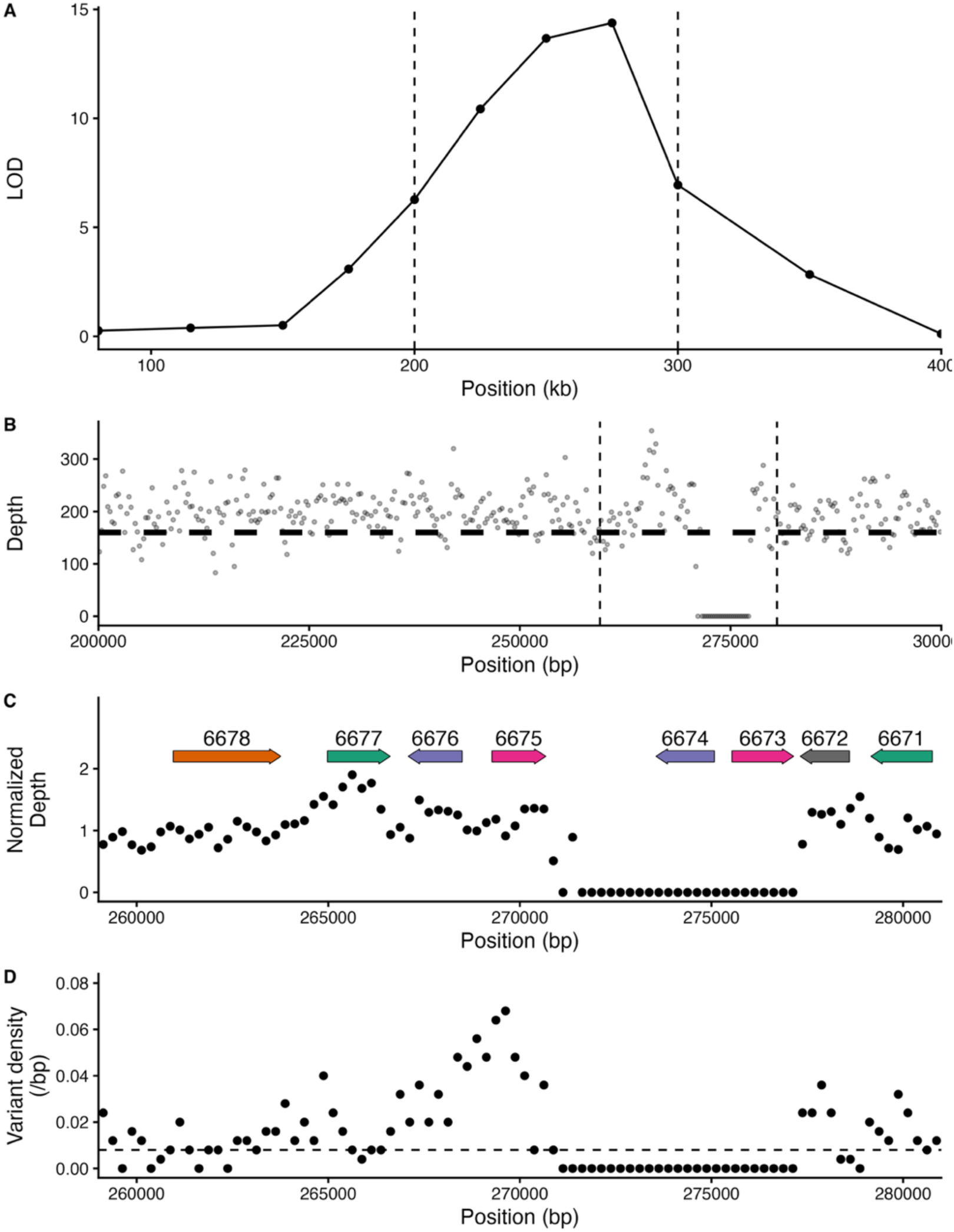
**Fine mapping of the *somA* locus**. A) LOD scores of QTL mapping using a panel of 47 recombinants across the 80kb – 400kb region of Chromosome 5 from the 8-24-15-7 inbred line. Dots indicate loci tested for allelic identity. B) Read depth of Java-6 mapped against the Okayama-7 reference genome. Dots indicate windows of 250 bp. C) Close-up of the region in B) indicating the genes surrounding the drop in coverage. For simplicity, gene names are shortened from full names (e.g. 6678 instead of CC1G_06678). D) Variant density across 250 bp regions, showing increased variant density near genes CC1G_06675 and CC1G_06676. Dots indicate average values across 250 bp windows. Horizontal dashed line indicates genome wide average of 0.008 variants per basepair.

**Figure 3:**
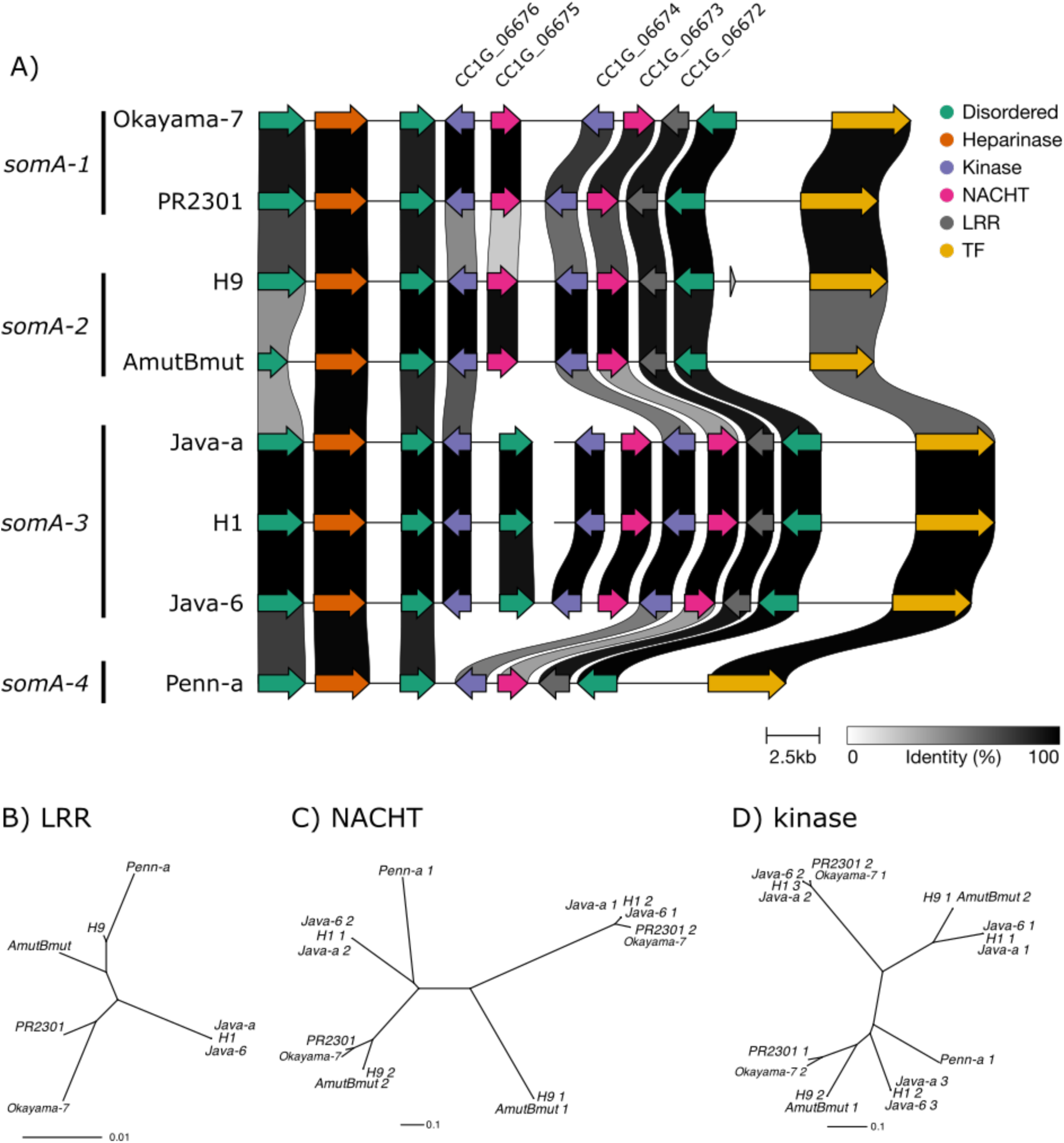
The *somA* locus is highly diverse within *C.* cinerea. A) Syntenic alignment of *somA* locus across *C. cinerea* strains. Each row represents the syntenic region of a particular strain, while arrows and direction indicate coding sequence orientation. Genes are colored according to similarity groups. Shaded regions connecting strains indicate amino acid sequence similarity, with amino acid similarity indicated in the text box. For Okayama-7, the 4 number code for the corresponding locus is given (i.e. 6675 = CC1G_06675). Note that for strains H1 and Java-a, the region is split across two contigs, indicated by the break in the genomic region. B-D) Neighbour joining trees of aligned amino acid sequences for LRR, NACHT, and kinase domains, respectively.

The region lacking sequencing coverage spanned the CC1G_06673 and CC1G_06674 genes (Fig. 2C). The other gene pair, CC1G_06675 and CC1G_06676, had notably higher genetic variation compared to the genome-wide average (Fig. 2D). In this region, three neighbouring genes, CC1G_06677 to CC1G_06675 had the most coding differences, both missense and synonymous variants (Table S1). However, the neighbouring two genes CC1G_06674 and CC1G_06673, while similar in size, had no variants annotated, neither missense nor synonymous. This pair was part of a repeated set, with CC1G_06676 and CC1G_06674 both predicted to encode kinases, and CC1G_06675 and CC1G_06673 predicted to encode NACHT domain proteins.

Due to the high sequence diversity at the *somA* locus between Okayama-7 and Java-6, and to further assess the variation at this locus across strains, we produced de novo assemblies of 7 additional *C. cinerea* samples from Illumina short read data, as well as hybrid assembly of Java-6 with long read data. These assemblies were significantly less contiguous than the reference Okayama-7 assembly, although the total genome assembly length was comparable (Table S1). Using conserved flanking genes we extracted the *somA* locus from each assembly. Across the seven genomes, we identified four highly diverse alleles, *somA-1* through *somA-4*. Between alleles, not only were individual genes diverse, with generally lower than 50% amino acid similarity, but the number of the kinases and NACHTs was highly variable. While the *somA* locus of each strain had at least a pair of one kinase and one NACHT domain (*somA-4*), some strains had two pairs (*somA-1*, *somA-2*), and some had two pairs and an additional kinase (*somA-3*). Across all strains, each NACHT gene was found in a tail-to-tail orientation with a kinase. Notably, directly adjacent to this region, there was a conserved leucine rich repeat gene (LRR). Between the Java6 and Okayama-7 parents of the cross, the LRR gene had more synonymous variants (20) compared to missense (6) (Table S2). The amino acid sequence similarity between alleles was high (∼99%) compared to the NACHT/kinase genes.

To determine if this gene arrangement was unique to the *somA* locus, we searched the Okayama-7 genome for domains similar to the LRR, NACHT, and kinase predicted protein sequences. This search resulted in a match to Chromosome 8, which also contained a NACHT and a kinase encoding gene, but in a head-to-head orientation, and with a LRR gene flanking on both ends (Fig. S2). Due to the similarity in protein domains and organization, we refer to this locus as the *somB* locus. Synteny analysis of the *somB* locus showed it was only present in Okayama-7 and AmutBmut, while in the other isolates the conserved flanking genes were present but the central three genes were absent (Fig. S1). Alignment of the two NACHT genes at the *somA* locus with the ones from *somB* (see below) revealed conserved NACHT diagnostic regions, primarily the glycine near the Mg2+ binding aspartate residue (Fig. S1).

The diverse sequences across alleles of the *somA* locus indicates that these alleles may be very old. To assess the conservation of this locus across wider mushroom-forming fungal taxa, we used the NACHT and kinase predicting coding sequences from both *somA* and *somB* to create custom hidden Markov models to search related taxa (Fig. 4A). Across related genera like *Coprinellus* and *Candellomyces* we found such genes in close proximity, also in head-to-head arrangement (Fig. 4B). Across these species, this ranged from 1-4 loci per individual, containing up to 10 NACHT or kinase domain proteins. Searches of more distantly related species, like the common button mushroom *Agaricus bisporus*, or the ectomycorrhizal tree-associated *Laccaria* failed to recover homologs of these genes in associated loci, as well as in associated proteins.

**Figure 4:**
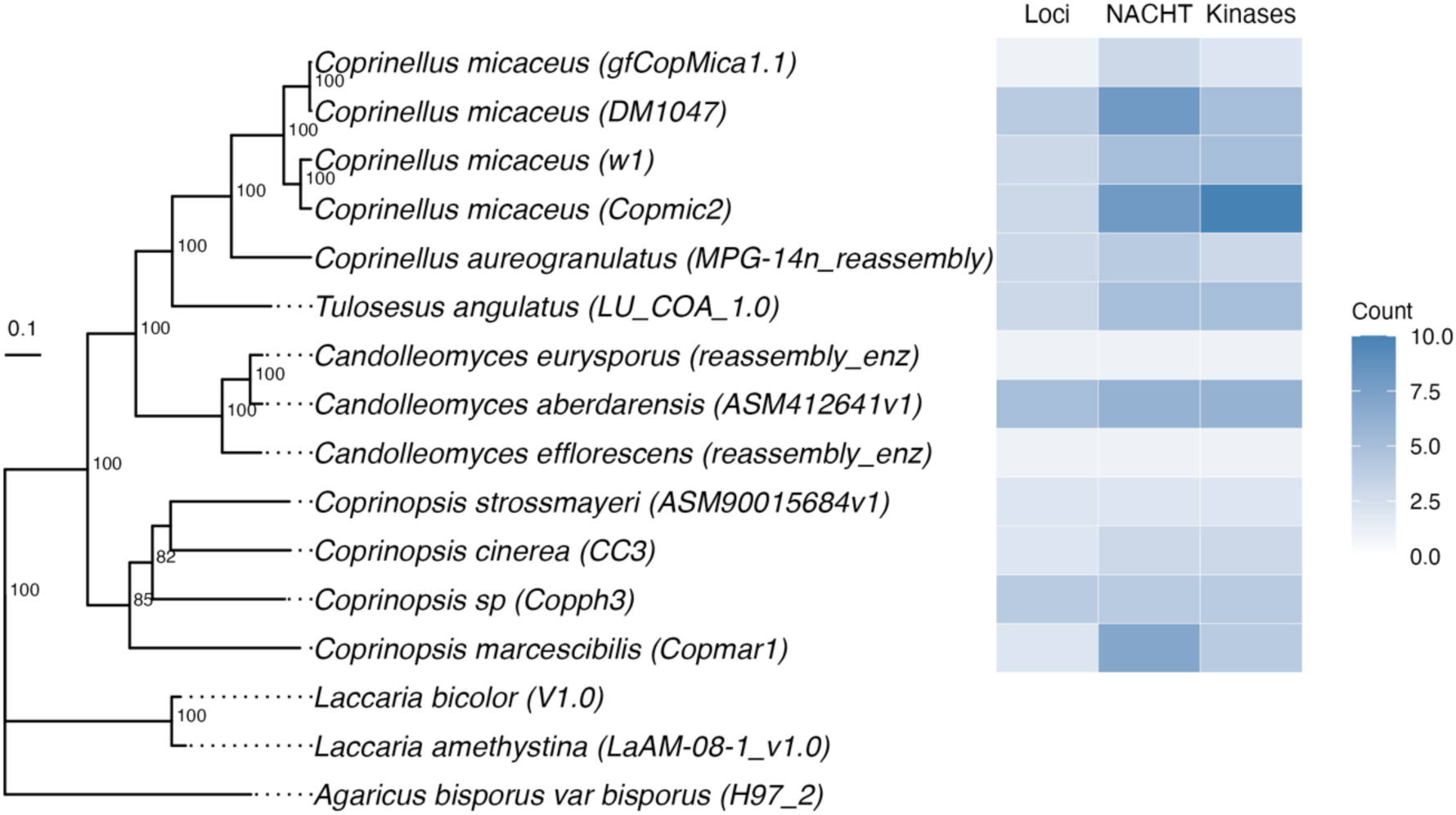
Association of NACHT and kinase genes in genomic regions across *Coprinopsis* and related species. To the left a maximum likelihood phylogeny based on the concatenation of three nuclear loci (RPB2, MCM7, and TSR1) with standard non-parametric bootstraps as branch support values. Branches are proportional to the scale bar (nucleotide substitutions per site). To the right, a heatmap shows the presence of putative loci as well as total numbers of NACHT and Kinase homologs identified based on a hidden Markov model trained on the *C. cinerea somA* alleles.

## Discussion

Here, we provide the first evidence linking a molecular mechanism to allorecognition in Basidiomycete fungi. In our F1 analysis we found that the ratio of offspring compatible with a parent was consistent with 3-4 segregating loci, similar to reports in other basidiomycetes. This is much lower than the ∼10-loci generally found in ascomycete species (Gonçalves & Glass, 2020; Nauta & Hoekstra, 1994). A trivial explanation for this reduced number of loci may be the smaller number of individuals studied compared to the larger ascomycetes studies (Anwar et al., 1993; Clavé et al., 2024; Garnjobst & Wilson, 1956). The loci we recover must segregate between the two selected parents, and other loci may be segregating in the species. However, this reduced number of loci could be biological, with a higher general allelic diversity for basidiomycete nonself recognition loci. For ascomycetes typically there are two, occasionally three, alleles for each locus while here we find four alleles from our eight monokaryotic strains. A higher number of alleles per locus means that a robust nonself recognition system would require fewer loci (Nauta & Hoekstra, 1994). Additionally, the dikaryon stage itself may be an explanation for why only 3-4 loci could be sufficient, with two genomes contributing combinatorial allelic diversity to the organism. It is unclear if these alleles act in a dominant/recessive or a co-dominant fashion. We have speculated on these interactions previously (Auxier et al., 2021),be an combinatorial but we lack experimental data to answer this point.

Although our segregation ratio indicated approximately 3-4 loci involved in SI, we recovered only a single QTL in the F1 offspring phenotyping, and the same single QTL in both backcrossed lines. We did recover a second locus located on chromosome 11 in one of the backcrossed lines, but since this was not found in the F1 compatibility testing, the relationship of this locus to incompatibility is unclear. A possible reason for consistent recovery of a single locus is that alleles at *somA* produce the strongest phenotype, and therefore largely determine the categorization of F1 offspring as compatible or incompatible. This would have a similar effect on the backcrosses, as they were screened for phenotypic incompatibility in each generation. Differences in the visible incompatibility phenotype between different loci has been observed in other species, like *Phellinus* (previously *Piptoporus*) (Adams et al., 1981; Rizzo et al., 1995). To identify other, potentially weaker loci, future work could use genetic markers targeting the *somA* locus to isolate incompatible lines which share the same allele for *somA*. A more technical explanation may be that an additional incompatibility locus resides in one of the reciprocal translocations found between Okayama-7 and Java-6 (Zolan et al., 1994).Okayama Many offspring were heterozygous for these regions, and these would complicate QTL mapping. Selection of more genetically similar parents may avoid this issue.

An important consideration to the results presented here is the use of a common nucleus. For all phenotyping experiments, strains were paired with the H1 nucleus to form dikaryons. Worrall termed this method a “common nucleus”, although this strategy had already been in use (5, 6). This method allows for genetic simplification of SI, since the common nucleus is fixed in all dikaryons, allowing the experimenter to control for its influence. However, simply because we kept this common nucleus constant, it should not then be inferred that this nucleus has no effect and is not epistatically interacting with the other nucleus. In fact, the opposite is likely true. As we have previously speculated, depending on whether these alleles are dominant or co-dominant, experiments utilizing different common nucleus partners may result in different genes identified (43). Based on the genome assemblies, within the cross used here, Java-6 and the common nucleus H1 have the same *somA* allele. Thus, the only difference between dikaryons at *somA* is the allele of Okayama-7. How the molecular interactions function within a dikaryon (i.e. the effect of a common nucleus that had a third allele) remains unclear. Future research in basidiomycete incompatibility would benefit from the use of additional common nuclei.

The general structure of *somA* alleles – a kinase that could function as a signalling molecule, a NACHT domain, and a LRR domain – is highly reminiscent of a general NLR-like genetic structure. The concept of NLRs in fungal nonself recognition has been extensively documented (Choi et al., 2012; Heller et al., 2018; Uehling et al., 2017). As well, the role of NLR-like genes in relation to immunity in both animals and plants is well established, and in fungi is being rapidly acknowledged as relevant. In the well-established models for the action of NLRs, a protein interaction domain, typically composed of repeated elements, like Ankyrin repeats, WD40 or Leucine-Rich Repeats, is thought to repress the action of the central NACHT nucleotide binding domain (Uehling et al., 2017). However, in *Coprinopsis* the protein domains of *somA* are found on independent proteins. Notably, the three components are encoded on opposite DNA strands, and so this cannot be a gene annotation artefact. In other NLR systems, it is thought that intramolecular forces within the multidomain NLR protein allow regulation (Heller et al., 2018). Understanding how this NLR-like protein functions in the absence of intramolecular interactions will be an exciting future study topic. If protein-protein interactions can be confirmed, this would also represent the first LRR type NLR found in fungi, as all other NLRs found in fungi use other repeat domains like Ankyrins or WD40 domains (Dyrka et al., 2014; Uehling et al., 2017). This organization of domains into individual proteins may explain why previous studies reported that LRR type NLRs are not found in fungi, as such searches begin with searching for proteins with multiple target domains (Dyrka et al., 2014). Additionally, the NACHT proteins we identify here are quite diverged from ascomycete NACHT proteins, hindering methods that extrapolate solely based on information from Ascomycete fungi.

The nonself recognition locus we identify here appears to be under strong balancing selection, with several highly divergent alleles found in a population. Particularly the NACHT and kinase genes from different alleles have little sequence similarity at either the nucleotide or protein level. Balancing selection is a common feature of nonself recognition systems, due to the increased fitness of individuals with rare alleles, resulting in alleles that can predate a species (J. Wu et al., 1998). In *somA*, the relatively high levels of synonymous variants compared to missense variants at the LRR gene indicate it is under purifying selection. The effect of balancing selection on nearby regions decreases with distance and recombination (Charlesworth, 2006). Theory predicts such regions should accumulate not only neutral mutations, but also deleterious mutations, although empirical studies are currently equivocal on this second point (Le Veve et al., 2023). Future work should focus on the wider biological consequences of these newly identified regions.

## Materials and Methods

### Culture Conditions

Strains were maintained on MYA (20 g/L Malt Extract, 15 g/L Agar) and grown at 37 °C for routine culturing. For sexual crosses, strains were co-inoculated onto test tubes with ∼2g of sterilized moistened horse dung (Moore & Pukkila, 1985), and incubated at 25 °C in 12h:12h day:night cycles. Basidiospores were collected by placing several gill pieces into water in microcentrifuge tubes, and heat shocked at 70 °C for 1 hour prior to dilution plating on MYA. Isolated colonies resulting from basidiospores were harvested using a tungsten needle (Moore & Pukkila, 1985). Long-term storage of strains was accomplished by freezing at-80 °C in 30% v/v glycerol.

For incompatibility screening, we phenotyped at the dikaryotic stage. For this, all strains were dikaryotised with homokaryon H1, with dikaryon formation being confirmed by visual inspection for clamp connections. For incompatibility testing, dikaryons were inoculated at opposite edges of either 9 cm petri dishes or in 100 mm 5×5 grid plates (Thermo Fisher Scientific Inc. Part #11339273). Incompatibility testing was performed on MYA, as initial comparisons with YPSS media (May, 1988) indicated little difference in the resulting phenotype.

### DNA Extraction

DNA for single isolates was isolated from mycelia scraped from agar plates using a CTAB buffer lysis step, followed by isopropanol precipitation, ethanol washing, and RNAse treatment (van de Peppel et al., 2021)(M. Nieuwenhuis et al., 2019). For in-silico bulk segregant analysis, mycelia were grown in liquid Malt Extract (20 g/L) and transferred to deep well 96-well plates. The mycelia were ground in a bead beater using 2 mm glass beads. DNA was extracted using a SPRI bead extraction, and DNA concentration was normalized using Quant-iT™ PicoGreen™ (ThermoFischer Scientific Inc.) fluorescence. Multiplex low-coverage barcoded libraries of offspring were performed using the hackflex protocol (Gaio et al., 2022). All DNA was sequenced using 150 bp paired-end reads on the Illumina Novaseq 6000 platform (Novogene Inc.).

### Single Isolate Genetic Analysis

DNA reads from single isolates were mapped to the reference Okayama-7 using bwa-mem (Li, 2013), and variants called with freebayes v1.3.1-19-g54bf409 (Garrison & Marth, 2012). Predicted effects of variants were annotated using snpEff v5.0e (Cingolani et al., 2012). For comparative genomics, genes were predicted using Augustus v2.5.5 (Stanke et al., 2006), and syntenic regions were compared using clinker (Gilchrist & Chooi, 2021).

For genetic analysis of offspring from Okayama-7 and Java-6, we used variants identified from Java-6 reads mapped to the Okayama-7 as a starting set. The low-coverage sequence of each offspring was mapped using bwa-mem as above, but only the known variants from Java-6 were used in downstream mapping analysis. For in silico BSA, we combined the low coverage sequences from offspring based on the compatible or incompatible phenotypes and genotyped the pools at the identified variant sites. We then used GATK VariantsToTable to format the input for use with QTLSeqR (Mansfeld & Grumet, 2018; Poplin et al., 2018). Genetic differences between bulks were assessed using the sliding window G’ value with a window size of 100 kb (Magwene et al., 2011).

### Comparative genomics

We collected published genome assemblies of species related to *C. cinerea* to look for homologs of *somA* (Table S2). To find candidate loci we searched for kinases and NACHT-containing genes that were close to each other, as defined below. For species that lacked a published genome annotation, we predicted *de novo* gene models using Augustus with default settings using the setting --species=coprinus_cinereus (Stanke et al., 2006).

To identify related kinase and NACHT domain-containing genes, we used MAFFT (Katoh & Standley, 2013) to align the kinases and NACHT gene from the *C. cinerea somA* and *somB* loci with default settings. This alignment was converted into a hidden Markov model using hmmerbuild and then each genome annotation was searched with hmmersearch (Finn et al., 2011). We only considered hmmer hits with an e-value smaller than 0.00001. For each hit, we extended the start and end flank coordinates of the gene model by 7 kb. We then intersected the location of kinases and NACHT-containing genes using BEDTools v. 2.31.1. The resulting intervals were further merged if they overlapped each other to determine the number of candidate loci.

To infer a species phylogeny, we used the *C. cinerea* sequence of the classic phylogenetics marker *rpb2* (Genbank accession number XM_001837441.2), as well as *mcm7* (XM_001833124.2) and *tsr1* (XP_001830678.2), two genes with good phylogenetic signal for fungi (Aguileta et al., 2008). We performed tBLASTn searches of the *C. cinerea* references on each species genome with the script query2haplotype.py v2.3 (available at https://github.com/SLAment/Genomics), using the following parameters: --task tblastn --haplo --minhaplo 200 --identity 60 --extrabp 1600. We aligned each gene with MAFFT v7.526 (Katoh & Standley, 2013) followed by manual curation using the CDS sequences as guides. The gene alignments were concatenated and given to IQ-TREE v 2.3.6 (Minh et al., n.d.) with automatic selection of substitution model (-m MFP) and 100 bootstrap pseudo-replicates (-b 100) to infer a maximum likelihood phylogeny. The tree and *somA*-loci counts were plotted in R with the libraries ape v5.8-1 (Paradis & Schliep, n.d.) and ggtree v3.10.0 (Xu et al., 2022). Rooting was done based on previous work (Wang et al., 2024).

## Acknowledgments

This work was supported by the Dutch Research Agency under grant ALWGR.2017.010. We thank Mayke Schelhaas for assistance with fine mapping of the *somA* locus. This work would not have been possible without the resources provided by the Fungal Genetics Stock Centre. We thank Shuru Mai for clarifying the strain names and relationships.

## Code and Data Availability

Code for genetic analysis and a Snakemake pipeline to locate *som-A* candidate loci is available in GitHub (https://github.com/BenAuxier/Coprinopsis_Somatic). Additional information, including *de novo* gene annotations are available at https://doi.org/10.5281/zenodo.20304762.

## Author Contributions

BA, AJMD, DKA conceived of research plan. BA, JB, KS, AvP, AJMD, DKA discussed research plans. BA, AJMD, DKA, JM designed genetic experiments. BA, JM analyzed genetic data. BA, JM performed laboratory experiments. LAV performed phylogenetic survey. BA wrote the initial draft of the manuscript. BA, AJMD, DKA, LAV edited manuscript. All authors approved final manuscript.

---

**Figure S1:**
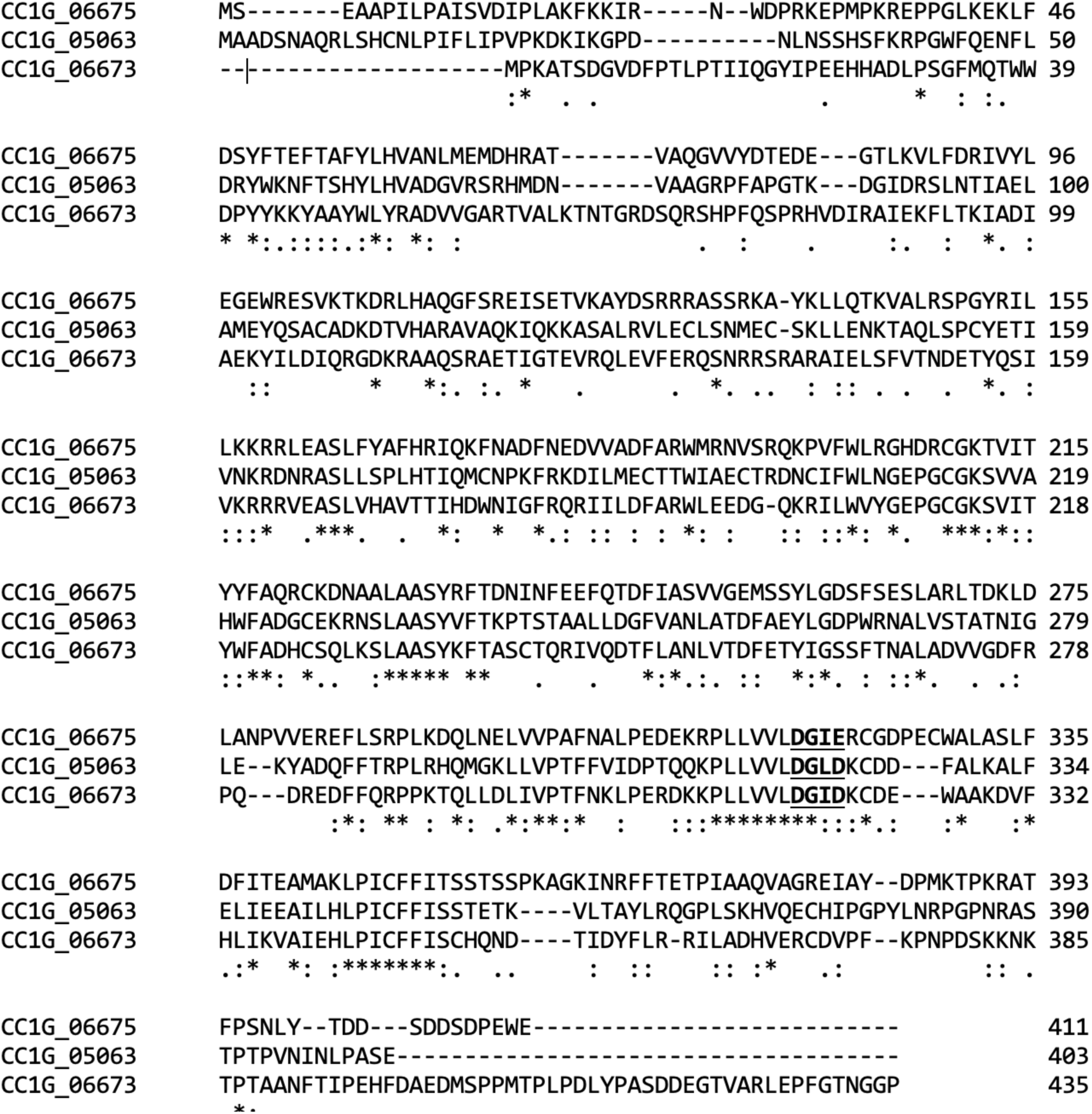
Multiple sequence alignment of the three NACHT protein sequences encoded by Okayama-7. Alignment produced using CLUSTAL-Omega. Asterisk below columns indicate complete conservation, while colons and periods indicate similar amino acid types. The defining residues surrounding the Walker B motif, mentioned in the main text, are underlined and bolded.

**Figure S2:**
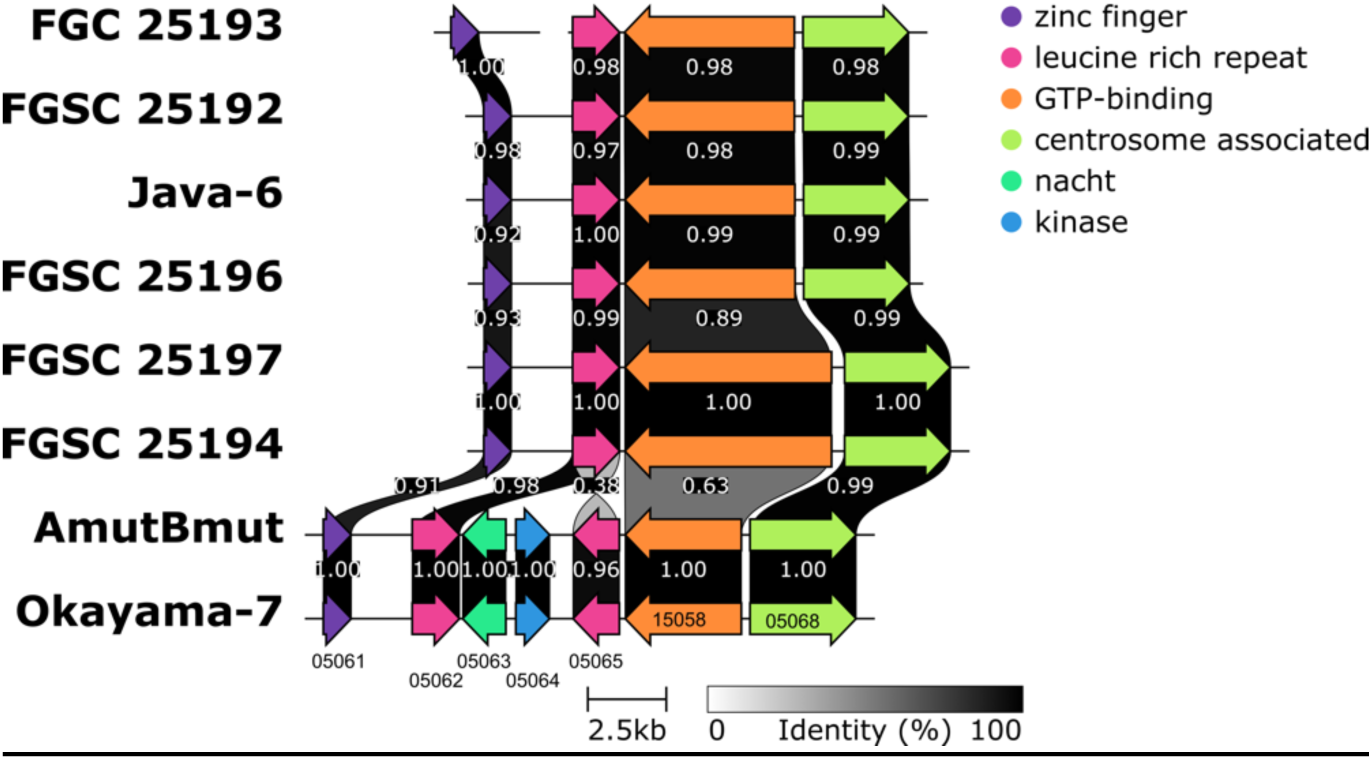
Synteny across *C. cinerea* strains across the *somB* locus on Chromosome 8 with similar protein organization as the *somA* locus. Genes and shading as in Figure 3.

**Table S1:**
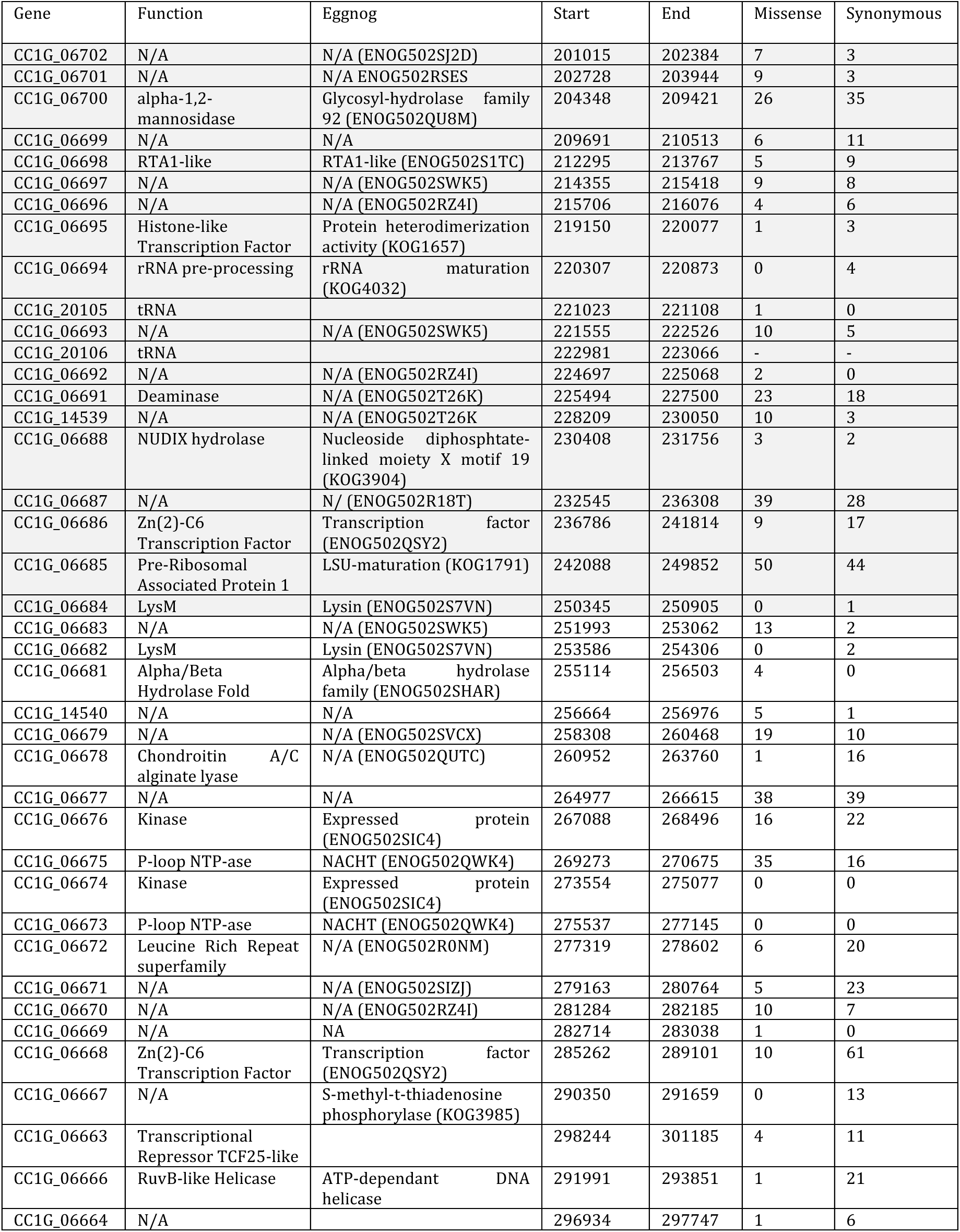
Predicted effects of Java6 coding variants across the *somA* locus. Genes in greyed section are found within the 200-250kb regions not indicated to be causal from the fine mapping.

**Table S2:**
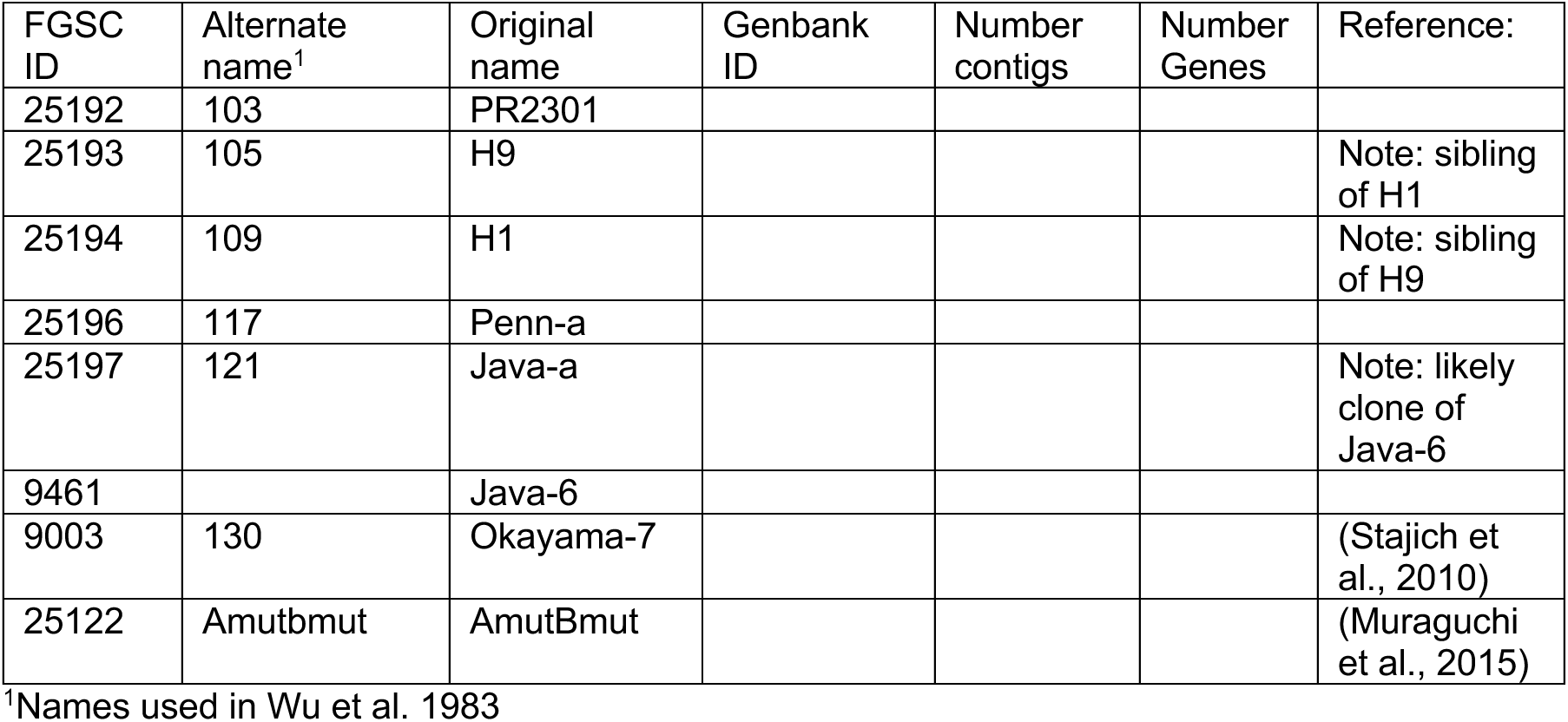
Statistics of *C. cinerea* genomes.

| FGSC ID | Alternate name<sup>1</sup> | Original name | Genbank ID | Number contigs | Number Genes | Reference: |
| --- | --- | --- | --- | --- | --- | --- |
| 25192 | 103 | PR2301 |  |  |  |  |
| 25193 | 105 | H9 |  |  |  | Note: sibling of H1 |
| 25194 | 109 | H1 |  |  |  | Note: sibling of H9 |
| 25196 | 117 | Penn-a |  |  |  |  |
| 25197 | 121 | Java-a |  |  |  | Note: likely clone of Java-6 |
| 9461 |  | Java-6 |  |  |  |  |
| 9003 | 130 | Okayama-7 |  |  |  | (Stajich et al., 2010) |
| 25122 | Amutbmut | AmutBmut |  |  |  | (Muraguchi et al., 2015) |

<sup>1</sup>Names used in Wu et al. 1983

